# Paired transcriptomic analyses of atheromatous and control vessels reveal novel autophagy and immunoregulatory genes in peripheral artery disease

**DOI:** 10.1101/2024.04.15.589664

**Authors:** Praveen Machiraju, Rajesh Srinivas, Ramraj Kannan, Robbie George, Stephane Heymans, Rupak Mukhopadhyay, Arkasubhra Ghosh

## Abstract

**Background:** Peripheral artery disease (PAD), a significant health burden worldwide, affects lower extremities due to atherosclerotic changes in the peripheral vessels. Although mechanisms of PAD have been studied, the molecular milieu of the plaques localized within peripheral arteries in patients are not well understood. Thus, to identify PAD lesion specific gene expression profiles precluding genetic, environmental and dietary biases, we studied the transcriptomic profile of plaque tissues normalized to non-plaque tissues from the same donors.

**Methods:** Transcriptomic analysis of 9 paired samples of PAD patients from south Indian population was performed with institutional ethics approval and written consent. Plaque tissues were histologically confirmed. Expression of a select target gene set was done by qPCR from the same tissues. Bioinformatic and network analyses were performed using various statistical tools.

**Results:** A total of 296 upregulated, 274 downregulated genes and 186 non-coding RNAs were identified. STAG1, SPCC3, FOXQ1 and E2F3 were key downregulated genes and CD93 was top upregulated gene. Autophagosome assembly, cellular response to UV, cytoskeletal organization, TCR signaling and phosphatase activity were key dysregulated pathways identified in the study. Telomerase regulation and autophagy were identified as novel interacting pathways using network analysis. Plaque tissue was predominantly composed of immune cells and dedifferentiated cell populations indicated by cell specific marker imputed gene expression analysis.

**Conclusion:** The current study identifies novel genes, non-coding RNAs, associated regulatory pathways and cell composition of the plaque tissue in PAD patients. The autophagy and immunoregulatory genes may drive novel mechanisms resulting in atheroma. These novel interacting networks and genes have the potential for PAD specific therapeutic applications.

## Introduction

Peripheral artery disease (PAD) is a progressive atherosclerotic disease affecting large peripheral arteries in humans often localized majorly to branch sites of the artery ^1^. PAD is one of the leading causes of morbidity due to atherosclerosis after coronary artery disease (CAD) and stroke ^2^. Prevalence of PAD increases considerably with age contributing to 1% of the total global deaths due to disease ^1,3^. PAD strongly correlates with occurrence of cardiovascular events thus contributing to an enormous economic burden in both developed and developing nations^4,5^.

Atherosclerotic lesions are complex, involving vascular endothelial and smooth muscle cells having dysregulated gene expression leading to interactions with monocytes and other immune cell types^6,7^. The altered secretory and behavioral properties of these cells along with accumulation of LDL in the intimal space leads to the plaque formation^8,9^. Although the sequence of atherosclerotic events has been described, the molecular pathophysiology of PAD has not been well elucidated. Recent transcriptional studies have identified immune and inflammation-related pathways to be majorly dysregulated, however these studies have either used age matched controls from different arterial beds or blood samples for transcriptomic studies ^10,11^. The literature on the transcriptomic studies of PAD is relatively limited compared to coronary artery disease or other types of vascular diseases. Even though both CAD and PAD are the manifestations of atherosclerosis in large arteries, significant differences in both presentation and molecular pathology exist between the two diseases making it imperative to further understand the molecular phenotype of atherosclerosis in PAD patients ^2^ ^12^ ^11^.

In this study we identified localized dysregulated genes which might contribute to pathogenesis and progression of atherosclerosis in peripheral artery disease and hence, we compared plaque material and non-atheromatous control vascular tissues from the same PAD patients. Our method of sample collection not only controlled for genetic variations but also nullified gender, risk factor exposure, diet and location-based differences in atherogenesis. This allowed for identification of genes and pathways which are directly involved and dysregulated in atherosclerotic plaques in patients. Further, a comprehensive transcriptomic analysis was performed to determine non-coding RNAs and cellular subtypes.

## Methods

### Sample collection and processing

20 pairs of non-diseased control samples and atherosclerotic plaque samples were collected from subjects of peripheral artery disease undergoing femoral artery bypass surgery at Narayana Institute of Vascular Sciences following informed consent and institutional guidelines (IRB Ethics approval no. NHH-MEC-CL-2015-355). The samples were immediately processed for RNA isolation and tissue fixation in formalin for staining. 9 samples out of 20 passed quality check and were used for further transcriptomic analysis. Cohort characteristics are listed in Table S1.

### RNA isolation and Quantitative real time PCR

Total RNA was isolated from control and plaque tissue using TRIZOL method according to manufacturer’s protocol (Invitrogen, Carlsbad, CA, USA). Briefly, the tissues were suspended in 500µl Trizol reagent, vortexed for 15 sec and incubated at room temperature for not more than 2 min. 200µl of chloroform was added for phase separation followed by centrifugation at 13200 rpm for 10 min. The upper aqueous layer was then added to a fresh tube containing 500µl of isopropyl alcohol and incubated for RNA precipitation at -40 ^0^C for 30 min. The precipitated RNA was centrifuged at 13200 rpm for 15 min, washed twice with 70% alcohol and suspended in RNAse free water. The concentration and purity of the extracted mRNA was assessed using spectrophotometer (Eppendorf spectrophotometer plus). cDNA conversion was performed using Bio-Rad iSCRIPT cDNA synthesis kit. Quantitative real time PCR was performed using SYBR green reagent (Kapa Biosystems Inc., USA). The quantitative real-time PCR (qRT-PCR) cycle included pre-incubation at 95°C for 3 min, 40 amplification cycles at 95°C for 10 sec, 58°C for 30 sec using a CFX Connect^TM^ real-time PCR detection system (Bio-Rad, Philadelphia, PA, USA). List of genes validated and corresponding primer sequences are given in Table S8.

### RNA Sequencing and Data analysis

RNA Sequencing was done using illumina HiSeq platform. Data analysis was carried out using standard bioinformatics pipeline for transcriptomic analysis. The analysis pipeline included trimming of the adapters and removing bases with low quality (Phred score > 30%), followed by contamination removal (rRNA, tRNA, mitochondrial sequences etc.) using Trimmoatic and Bowtie (version-2.2.4). The pre-processed reads were then aligned to human genome (hg19) using HISAT2. The reads aligned were used for finding differential gene expression (DGE) using Feature Counts and DeSeq2.

### Differential gene expression calculation

The raw read counts for control and case samples were normalized using DESeq2. log2 (foldchange) values were found to be normally distributed. From this distribution, the genes which were found to be 2 standard deviations away from the mean (mean±2SD) were considered as differentially expressed. The downstream annotation was done using these differentially expressed genes. The Gene Ontology annotations were obtained using AmiGO2. Amigo2 performs GO enrichment analysis for the given up or down regulated gene set. Based on that information it will categorize the gene set as over-represented (+) or under-represented (-). It also includes fold enrichment information for differential expressed genes and Bonferroni corrected p-values.

### Tissue processing and staining

Tissue sample from patients was immediately fixed in formalin. Following fixing, the tissues were embedded in paraffin wax and 4µm sections were taken using a Leica microtome 2235. The sections were kept on a hot plate for 1 hour at 65°C and processed for hematoxylin and eosin staining (H& E) staining. For H&E staining, the tissue was hydrated using xylene and alcohol for 5 min each. Tissues were stained with Harris’s hematoxylin for 10 min followed by washes in acid alcohol and 2% sodium bicarbonate for 2 min each. Eosin staining was done for 1min. The tissues were hydrated by washing in alcohol and xylene for 2 min each and mounted using a DPX mounting medium.

### Bio-informatic analysis and visualization

Data visualization was done using orange ^13^and gene network analysis was done using KEGG pathway analysis. miRNA identification and differential expression quantification of non-coding RNA was performed using miR Master 2.0 ^14^. Cell marker 2.0 was used for cell imputations ^15^. Cell enrichment scores were obtained from dividing number of identified gene markers with number of gene markers available in the database. Gene networks and gene enrichment network visualization was performed in Cytoscape (version3.10.1) using ClueGO^16,17^.Transcription factor enrichment analysis was performed using ChEA3^18^.

## Results

### Transcriptomic profiling reveals novel dysregulated genes in PAD

To validate our sample collection method (Figure1A), we performed H&E staining to confirm that the controls used in this study were non atherosclerotic. Dense eosin staining of plaque in the atherosclerotic tissue sample was observed but not in controls (Figure1B). In addition, the gene expression of atherosclerotic markers such as Intracellular adhesion molecule-1(ICAM), Vascular cell adhesion molecule-1 (VCAM) and Monocyte chemoattractant protein-1(MCP-1) were assessed using qRT-PCR. ICAM1 (FC=9.23, p<0.05), VCAM1 (FC= 2.45, p=0.06) showed significant upregulation and MCP-1 showed 1.7-fold higher expression in atherosclerotic tissues confirming that the molecular expression patterns between the matched disease and control tissues from patients were identified correctly (Figure1C).

**Figure 1:**
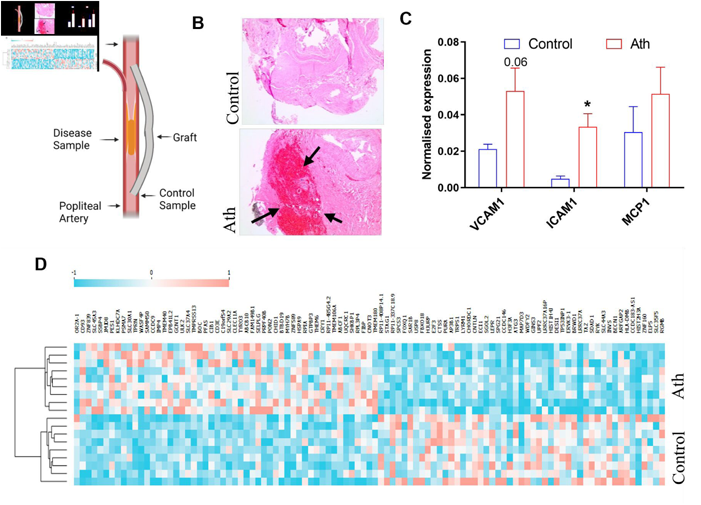
RNA Sequencing from atheromatous tissues obtained from subjects undergoing femoro-popliteal bypass surgery show altered gene expression profiles. (A) Schematic shows the site of sample collection. (B) H & E staining showing atheromatous tissue as compared to control (Ath-Atheromatous plaque sample). Arrows indicate plaque region of the sample marked by dense eosin staining (C) Normalised gene expression profiles of ICAM1, VCAM1 and MCP1 in atheromatous tissue (Ath) and control tissue n=3, *p<0.05, *Mann Whitney U test*. (D) Heatmap of top differentially regulated genes identified through RNA sequencing. Each column of the heatmap represents one sample. Genes are clustered using k means clustering.

A total of 296 significantly downregulated and 274 upregulated genes were identified in combined RNA sequencing analysis of 9 paired samples. Figure 1D shows the heatmap of top differentially regulated genes identified in this study. Top downregulated genes were STAG1(log2FC=-0.63, p<0.05), SPCS3(log2FC=-0.62, p<0.05), SAR1B (log2FC=-0.61, p<0.05) and USP8(log2FC=-0.61, p<0.05). Mutations in STAG1, (Stromal antigen1 cohesion subunit 1) have been reported to predispose children with hematological malignancies and it is important to maintain genomic stability^19^. Downregulation of STAG1 in PAD would indicate loss of genomic stability and predisposing the genome to DNA damage. USP8 is important for endosomal trafficking in atherosclerosis and is induced in presence of growth arrest during cell-cell contact and is also important in RAS signaling and Wnt pathway regulation^20^. Top downregulated transcription factors comprised of FOXQ1(log2FC=-0.61, p<0.05) and E2F3 (log2FC=-0.61, p<0.05). E2F3 has been implicated in atherosclerosis, ^21^ but the role of FOXQ1 atherosclerosis is unknown.

Top upregulated genes identified included immune related and cell surface adhesion genes like CD93 (log2FC=0.76, p<0.05), JMJD8(log2FC=0.6, p<0.05) and PES1(log2FC=0.61, p<0.05). zinc finger protein encoding gene ZNF839(log2FC=0.72, p<0.05), solute receptor family member SLC45A3(log2FC=0.71, p<0.05) and SSBP4(log2FC=0.68, p<0.05). CD93 is a well-known transmembrane receptor and is important in promoting monocyte adhesion and further aiding in macrophage migration^22^. Table S2 lists top dysregulated genes identified through transcriptomic analysis. Gene expression analysis using quantitative real time PCR validated findings from RNA seq analysis. Genes such as AP3B1, LYRM1, USP8 and Rab11 were found to be downregulated in transcriptomic data as well as gene expression analysis in the plaque tissues. Whereas, genes such as FGFR3, LOXL1 and ULK2 were found to be upregulated. (Figure S1).

### Gene ontology analysis identifies key pathways in PAD

Key downregulated biological processes comprised of autophagosome assembly (FE=6.97, p>0.05), vesicle mediated transport (FE=4.23, p>0.05), cellular response to DNA damage (FE=2.74, p>0.05) (Figure 2A, Table S4). Autophagosome assembly and vesicle transport included genes such as BECN1, ATG3, RAB11A, UBQLN1 and SAR1B. Upregulated biological processes included enrichment of actin cytoskeleton organization (FE=4.18, p>0.05) and actin filament polymerization (FE=8.9, p>0.05) (Figure 2B). The pathways identified are indicative of multiple dysregulated pathways related to primarily endothelial cells and macrophages, both crucial drivers of disease progression. Upregulated pathways also included focal adhesion genes, MAPK signaling and TCR signaling (Figure S2B, Table S3). Downregulated molecular functions included pre-mRNA intronic binding (FE=27.5, p>0.05) and R-SMAD binding (FE=10.48, p>0.05) (Figure 2C), and upregulated molecular functions were DNA photolyase (FE=76, p>0.05), DNA helicase activity (FE=76, p>0.05) etc, (Figure 2D, Table S3). Overall, the gene ontology from upregulated genes indicated a pro-inflammatory milieu and significantly higher actin polymerization. Pathway enrichment network analyses revealed novel pathway interactions such as telomerase regulation and autophagy regulation (Figure 3A). Pathways identified using upregulated genes were found to be independent of each other (Figure 3B). Figure 3C and Figure 3D show differentially expressed genes in pathways identified.

**Figure 2:**
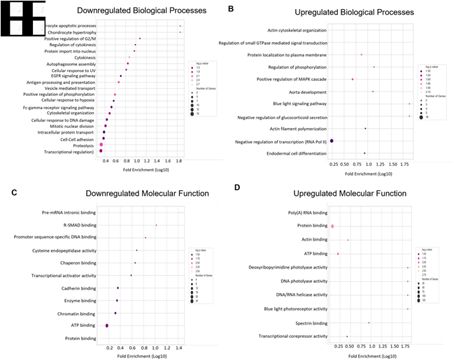
Altered biological processes associated with PAD. (A) Downregulated biological processes in PAD. (B) Upregulated biological processes (C) Downregulated molecular functions and (D) Upregulated molecular functions associated with PAD. Colour scale in bubble plots indicates log p-value, size of the bubble denotes number of genes represented and X-axis represents log10 value of fold enrichment.

**Figure 3:**
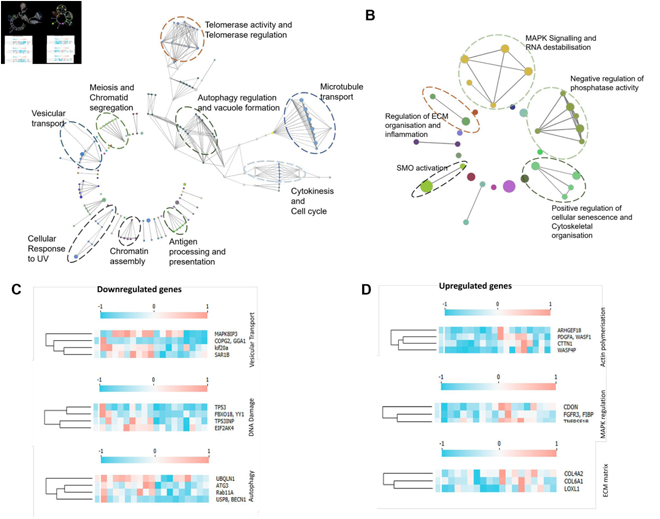
Network analysis identifies unique pathway interactions in PAD. (A) Downregulated pathway networks in PAD, encircled network interactions represent novel interactions found in PAD. (B) Upregulated pathway networks in PAD, encircled network interactions represent novel pathways identified in PAD. (C) Heatmaps show relative fold change of downregulated genes specific to identified pathways from the dataset. (D) Heatmaps show relative fold change of upregulated genes specific to identified pathways from the dataset. Columns indicate genes and rows indicate independent samples. For heatmaps gene expression is normalised with average gene expression value and genes with similar k mean are clustered together.

### Gene networks involved in PAD pathogenesis reveal novel interactions

The search tool for retrieval of interacting genes (STRING) (https://string-db.org) was used to construct gene networks. Cytoscape software version 3.10.0 was used to visualize the networks. Upregulated gene network analysis had 86 nodes and 91 edges with 2116 average number of neighbors and downregulated gene network analysis had 196 nodes, 452 edges and 4612 average number of neighbors. Gene network analysis of top 100 downregulated genes identified novel interactions among genes such as CD4 and Beclin1 as well as CTSS, which is known to be impaired in atherosclerosis^23^. H4C6, a histone coding gene interacted with STAG1, KIF20A and other histone protein coding genes such as H3-3B (Figure 4A). Solute carrier family genes such as SLAC30A1, TMC6 and SLAC45A3 clustered together in the upregulated gene network analysis. Mitochondrial and metabolism related genes such as NDUFS7, TMEM126B and EIF3G were clustered together indicating a metabolic impairment in atherosclerosis (Figure 4B). Together, gene interaction network analysis of downregulated genes predicted novel networks relating to autophagy and T-lymphocyte function as well as interactions among histone protein coding genes and genes related to DNA stability. Novel interactions of mitochondrial genes with cytoskeletal genes such as PARVB were observed in the downregulated set. Downregulated TFs (transcription factors) enriched in the dataset included CREB1, GABPA and TCF12 (Figure 5A) and upregulated TF network included ZNF316 and FOXP4 (Figure 5C). Most of the downregulated transcription factors in the network pertained to immune response and lipid biosynthesis (Figure 5B) while bioprocesses like negative regulation of transcription, DNA transcription and RNA3’ processing were upregulated (Figure 5D).

**Figure 4:**
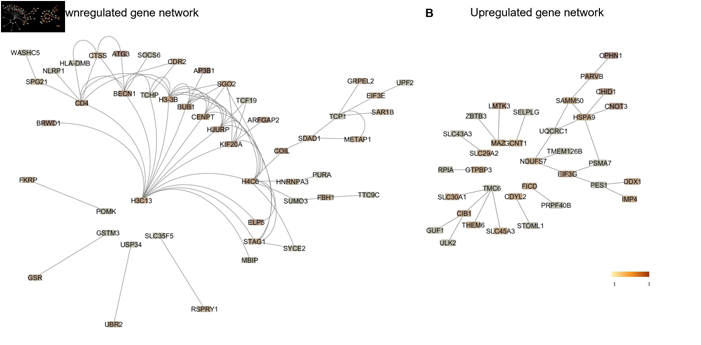
Gene network interactions in PAD. (A) Downregulated gene network of top 100 downregulated genes in PAD showing H3C13 having most interactions. (B) Upregulated gene network of top 100 upregulated genes in PAD. Gene network interactions were studied using STRING and visualized using Cytoscape. Color Scale represents shortest average pathlength.

**Figure 5:**
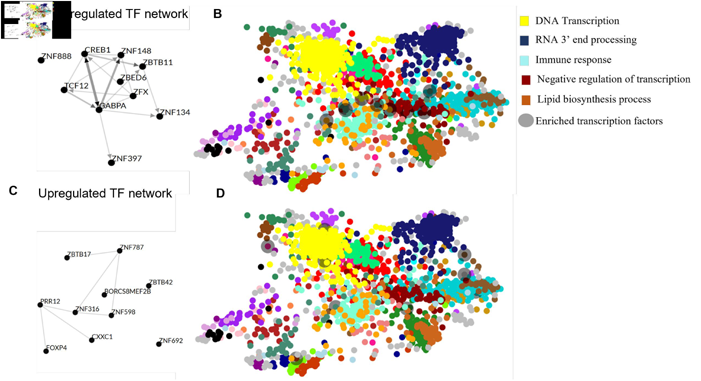
Novel transcription factor networks in PAD. (A) Transcription factor network analysis of top 10 ranked transcription factors from downregulated genes identified in plaque samples compared to controls. (B) Downregulated transcription factors overlapped to colour maps obtained from Genotype-Tissue Expression TF network (GTEx TF) clustered according to GO enrichment. (C) Transcription factor network analysis of top 10 ranked transcription factors from upregulated genes identified in plaque samples compared to controls as analysed using ChEA3. (D) Upregulated transcription factors overlapped to colour maps obtained from Genotype-Tissue Expression TF network (GTEx TF) clustered according to GO enrichment.

### Non-coding RNA analysis identified novel non coding-RNAs in PAD

Four types of regulatory ncRNAs namely, long non coding RNA (lncRNA), circular RNA (circRNA), miscellaneous RNA (miscRNA), piwi interacting RNA (piRNA)were mapped using in the dataset using miR Master v2.0.A total of 118 dysregulated lncRNA, 44 dysregulated circRNA, 17 dysregulated miscRNA and 7 dysregulated piRNA were identified (Figure 6A). We could not detect significantly differentially expressed miRNA since only 0.001% of sequences could be mapped to existing miRNA database (Figure 6B). Most of the ncRNA identified in the study were found to be novel. CircRNA targeting MACF1 (Log2FC=1.36) loci, MEF2A (Log2FC=1.2) loci were upregulated whereas PSEN2 (Log2FC=-1.47) and WDR67 (Log2FC=-1.42) loci circRNA were downregulated. MACF1 and MEF2A loci circ RNA indicates an upregulation in actin polymerization and inflammation as exhibited in the gene ontology identifications. AL117329.1(Log2FC=1.3, p<0.05) and AL732437.2 (Log2FC=-1.8, p<0.05) were top upregulated and downregulated lncRNAs respectively (Figure 6C) (Table S5).

**Figure 6:**
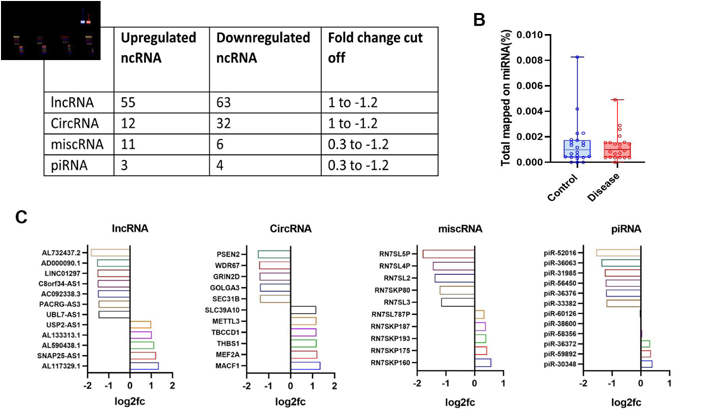
Non-coding RNAs in PAD provide insights to underlying dysregulated mechanisms. (A) Table shows number of upregulated and downregulated non-coding RNAs identified along with respective fold change cut off values used in the study. (B) Percentage of sequences from RNA-sequencing mapped to miRNA database. (C) Bar plots showing differential expression of non-coding RNAs as identified using miR Master v2.0 in PAD cohort.

### Cell profiling identified key cell types involved in PAD

To understand the cellular subtypes that differentially regulated genes could be attributed to in PAD, we imputed cell fractions in PAD from the list of upregulated and downregulated genes using the CellMarker 2.0 database. Figure 7A shows downregulated genes mapped to different cell types and Figure7D shows upregulated genes mapped to specific cell types. A total of 59 contributing cell types were identified within the downregulated gene list (Figure 7C) and 56 cell types in the upregulated gene list (Figure 7F) (Table S6). The cell types were then divided into seven categories depending on their biological function. Stromal cell markers were enriched with a gene enrichment of 0.033 in both upregulated and downregulated gene lists (Figure B, E). Immune cell markers were the highest mapped genes markers in both upregulated and downregulated genes, 183 and 234 respectively. (Table S7).

**Figure 7:**
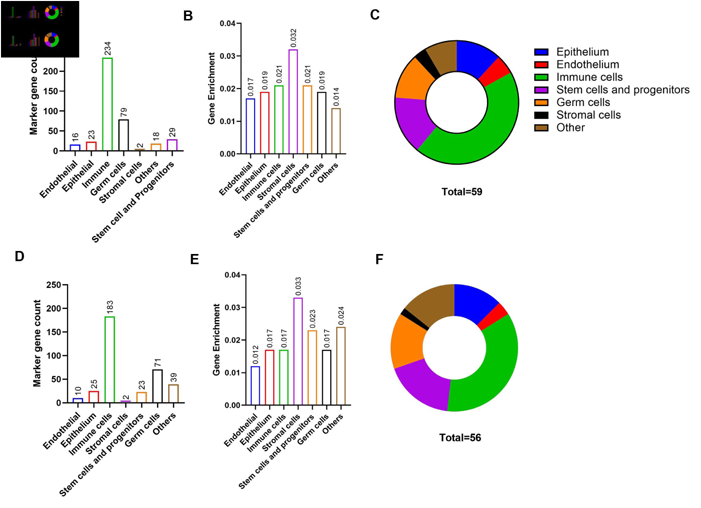
Cellular composition of PAD plaques. (A) Marker gene counts (B) Gene enrichment and (C) Major cell types identified and their proportions imputed using downregulated genes. (D) Marker gene counts, (E) Gene enrichment and (F) Major cell types identified and their proportions imputed using upregulated genes. Gene enrichment was calculated as a ratio of number genes mapped to specific cell type and total number of cell specific genes.

## Discussion

Comparative molecular studies to understand the disease complex disease phenotypes typically involve healthy donors and patients where a critical limitation is that the basal expression levels of genes and the genetic makeup between controls and patients do not always match. Therefore, transcriptomic analysis of atheromatous peripheral artery was compared to a matched non-affected patent artery sample of the same patient. The genes identified in such paired analysis from patients can be held in greater confidence for their involvement in disease progression as the study design negates the identification of those genes which might be differentially expressed elsewhere in the vasculature due to age-related, genetic, metabolic or co-morbid conditions of the patients affecting broader changes in physiology. The study not only provides a novel approach to better identify disease related genes but also is one the first reports from a south Indian patient cohort where molecular expression phenotypes were not investigated previously.

Differentially expressed genes such as CD93, SAR1B and USP8 detected using such paired comparisons are novel discoveries in PAD. SAR1B functions as a transmembrane receptor for lipids and is important for maintaining lipid homeostasis. Mutations in SAR1B may cause chylomicron retention disease ^24,25^. FOXQ1 is a transcription factor involved in invasion and metastasis through EGF receptor pathway ^26^. Although many downregulated genes are relevant to atherosclerosis, their role in molecular mechanisms of disease need further studies. CD93, which was upregulated in PAD atheroma, is a cell surface lectin receptor expressed on macrophage and endothelial membranes. CD93 is crucial for intra cellular adhesion and clearance of apoptotic cells ^27^. In endothelial cells, CD93 promotes cell-cell adhesion through beta integrin activation and fibrillogenesis leading angiogenesis^28^. SLC45A3, which was also elevated in atheroma, plays an important role lipid metabolism ^29^. Gene expression studies in PAD have identified genes like MMP9, MMP12, SPP1 and APOD however, these identifications compare gene expressions in plaques from different arterial beds and not from the same subject^11^. The pathways identified in previous studies also predominantly are of immune and inflammation related pathways^30^. Other studies either targeted specific cells such as macrophages or the identifications were not from plaque material of PAD subjects^31-33^. Since the dysregulated genes specific to the plaque milieu of the PAD patients are controlled to their own patent vessel tissues, many of the genes that have been identified in other studies did not show significance in our study. Such genes may be altered in the non-lesion areas of the vessels or be associated with more global changes occurring within the PAD patient. Therefore, this analysis furthers our understanding of advanced PAD and identifies these genes as specific targets for future therapies directed towards the atheromatous lesions in advanced disease.

Pathway enrichment analysis primarily identified vacuole assembly, autophagy and EGFR signaling as key downregulated pathways apart from cellular response to UV and protein localization. TCR-signaling, IL-17 signaling, ECM regulation and phosphatase activity are upregulated in pathway network analysis. In agreement with previous findings, proteinase and peptidases expression are upregulated in femoral artery plaques ^34^, phosphatase activity and related networks are upregulated, and regulation of MAPK activity by phosphatases increases foam cell formation and VSMC migration during atherosclerosis^35^. Interaction between telomerase activity and autophagy related processes was a novel finding from our study. Rescuing autophagy has been shown to ameliorate atherosclerosis in mice^36^. Although reduction in telomerase reverse transcriptase (TERT) expression levels and telomerase activity have been implicated in atherosclerosis, definitive mechanisms have not been elucidated^37^. Our findings suggests that telomerase regulation and autophagy processes together might confer an antiatherogenic effect in PAD. Moreover, pathway interaction networks also indicate a possible regulatory mechanism among genes involved in these processes.

Non coding regulatory RNA (ncRNA) such as long non coding RNA (lncRNA), circular RNA (circRNA), miscellaneous RNA (miscRNA), piwi interacting RNA (piRNA) and microRNA (miRNA) play crucial roles in expression and regulation of genes and proteins through ncRNA-DNA, ncRNA-mRNA, ncRNA-protein and ncRNA-ncRNA interactions^38,39^. Studies to identify miRNA have focused on circulating miRNAs using whole blood or PBMCs ^40,41^ or studying specific PAD related events in mice models ^42^ but data in human subjects have not yet been reported. LINC01297, UBL-7 AS1 and SNAP25 AS1 are novel lncRNAs identified in this study, which are known to regulate infiltration of immune cells, cell proliferation and vesicular transport^43,44^. The circRNA PSEN2 regulates PSEN2 gene which is involved in cleaving late endosomal substrates was dysregulated in atheromatous tissues^45^. Together, non-coding RNA identification indicated autophagosome disassembly, upregulation of microtubule assembly and deregulation in immune modulation. Although we could identify miRNAs from our data, no statistically significant differentially expressed miRNAs were observed which may be due to the study design.

Classification of the cell types showed that although immune cell types were largely represented, it was the stromal cell gene signatures which were enriched in femoral artery plaques. These findings support the notion that femoral artery plaques are much more stable owing to denser stromal matrix. However, cell subtype identification analysis may be limited by the available databases and cell specific marker information on atherosclerosis.

In conclusion, by using paired disease-control analysis, our study uncovers key genes and interacting networks that are signatures of plaques in peripheral arteries of PAD patients. Further functional studies of the genes identified may lead to the development of targeted therapies for peripheral artery disease.

## Acknowledgements

We gratefully acknowledge the support from Dr. K. Bhujang Shetty and Dr. Rohit Shetty for their support and providing resources during the experimental studies. We also acknowledge Dr. Ramesh Tripathi for his support and advice on the clinical aspects of the study.

## Supplementary Materials

Table S1: Study cohort characteristics

Table S2: Dysregulated genes in PAD

Table S3: Gene Ontology of upregulated genes in PAD

Table S4: Gene Ontology of downregulated genes in PAD

Table S5: Differentially regulated non-coding RNAs identified in PAD

Table S6 – S7: Cell imputations and marker gene counts table

Table S8: Primers sequences of genes used for data validation

Figure S1: Gene expression analysis for validation

Figure S2: Gene Ontology analysis of dysregulated genes

